# Continental scale genetic diversity supports geo-clustered ecoregions connectivity

**DOI:** 10.64898/2026.11.05.764579

**Authors:** A. C. Riley, M. Hajibabaei

## Abstract

Connectivity is fundamental to ensuring that ecosystems can recover after impact ^1–3^. The movement of organisms between regions can occur at the individual level or over multiple generations ^4^. Genetic variation among species in different locations has important implications for population status and health ^5,6^. Here we utilize large-scale DNA metabarcoding data of benthic arthropods to study continental scale connectivity and gene flow. We compared structural connectivity, represented by an adjacency matrix of ecoregions, and functional connectivity at various levels of organization, represented by the β-diversity of exact sequence variants and species, between ecoregions. While structural and functional connectivity were correlated, there was a distinct divide between eastern and western ecoregions that approximately aligned with the Rocky Mountains. Within the western ecoregions, species β-diversity was not correlated with ecoregion adjacency however exact sequence variant β-diversity was correlated. Additionally, species had higher intraspecific genetic differentiation in the western ecoregions than the eastern ecoregions. This implies that in the western ecoregions gene flow is more restricted. Species in areas where gene flow is limited will be more vulnerable to impact and have slower recovery. Understanding where gene flow is constrained is important to conservation planning.

**Significance Statement:** Maintaining and improving ecological connectivity is essential to the recovery and resilience of organisms to climate and land use changes. Connectivity is driven by organism movement, movement shapes gene flow, and gene flow can be identified by observing genetic structure. We demonstrate that DNA metabarcoding data can be used to observe genetic structure in samples that span the United States. The genetic structuring we observed suggested that there is less gene flow in the Rocky Mountains indicating that these population are more isolated. By identifying where connectivity is weak, conservation planning can be focused on protecting these vulnerable regions.

## Main

Climate change and habitat loss are environmental drivers occurring at a global scale. While these drivers are occurring at broad-scales, the response is unique to local geographic regions. Understanding how and why responses are different is essential to developing action plans for conservation, sustainable development and environmental stewardship.

Biodiversity in freshwater ecosystems is in decline ^7^. Current conservation strategies in the United States are not sufficient to mitigate loss and recovery in European systems has stalled ^8,9^. Ecological biomonitoring studies rely on bioindicator species such as benthic macroinvertebrates to understand ecosystem status because these organisms are highly sensitive to environmental perturbations ^10^. However, effective monitoring is hamstrung due to a lack of fine-scale taxonomic information leading to knowledge gaps or inconsistency in observing change especially at larger scales ^11–13^.

Connectivity plays a key role in species resilience and recovery. If an impacted area has a population that is reduced or lost, a connected population can repopulate that area introducing lost genetic or functional diversity ^1–3^. Thus, understanding how populations are connected can aid in conservation planning by determining vulnerable species and regions that need to be prioritized.

In ecology, connectivity has been defined in various ways, but it has two well defined components ^4,14–16^. Structural connectivity, which is defined by patterns in space, and functional connectivity, which is defined by how organism movement is affected by patterns in space. Using a river as an example, structurally, it can be seen as dividing land or it can be seen as a network linking bodies of water. Functionally, a terrestrial organism may have its movement restricted by the river where it acts as a barrier, while an aquatic organism may be limited to only moving through the river. Understanding the interaction between these two components is essential to understanding how spatial patterns shape the movement of organism and the flow of genetic information.

In landscape ecology, the strictest definition of functional connectivity is the direct observation of individual organism movement, however measuring direct movement can be challenging depending on the organism of study ^16,17^. Changes in organism composition (β-diversity) has been used as proxy for functional connectivity ^2,18^. If two locations share the same organisms, at some point in time an organism had to travel between the locations. This could have occurred by the direct movement of individuals or over multiple generations ^4^. However, species distribution has been suggested to be only linked to functional connectivity and not represent actual functional connectivity ^16^, this is because it only provides information on whether or not there are populations of a species present at those locations. Populations could be isolated from each other, meaning there will be no gene flow between them. Therefore, measuring gene flow, in terms of timescale, is closer to connectivity by movement than β-diversity ^4^.

Using reference databases, DNA metabarcoding can provide species-level information that can be used to track species distributions. However, DNA metabarcoding also provides critically underutilized genetic information that falls somewhere between the individual and its taxonomic classification ^19^. Where this falls exactly will depend on the organism being studied, but there is support for using exact sequence variants (ESVs) as proxies for haplotypes which can be treated as genetically distinct groups ^5,6^. Understanding the genetic structure of ESVs within a species could provide insights into connectivity through gene flow.

In this study, we used 19 ecoregions (also referred to as ecoclimatic domains) to examine connectivity on a United States continental scale ^20,21^. We define structural connectivity on the spatial arrangement of ecoregions. We define functional connectivity as the β-diversity between ecoregions, where β-diversity is derived from genetic data from DNA metabarcoding analysis of aquatic arthropods collected as a part of the U.S National Science Foundation’s National Ecological Observatory Network (NEON). We compare spatial autocorrelation between structural and functional connectivity and use percolation thresholds to investigate weighted ecoregion graphs based on functional connectivity. Finally, we use intraspecific genetic differentiation to investigate patterns in gene flow.

### Spatial autocorrelation in the ecoregion adjacency matrix

The relative position of ecoregions was represented by a graph, where an edge forms between two nodes (ecoregions) if the borders between two ecoregions is shared (Fig. 1a). This graph can be expressed as a matrix, where pairs of ecoregions are either adjacent, represented by 0 in the matrix, or not adjacent, represented by 1. By comparing the β-diversity dissimilarity matrix of ecoregions to their adjacency matrix, the functional and structural relationship between ecoregions was compared.

**Fig. 1.**
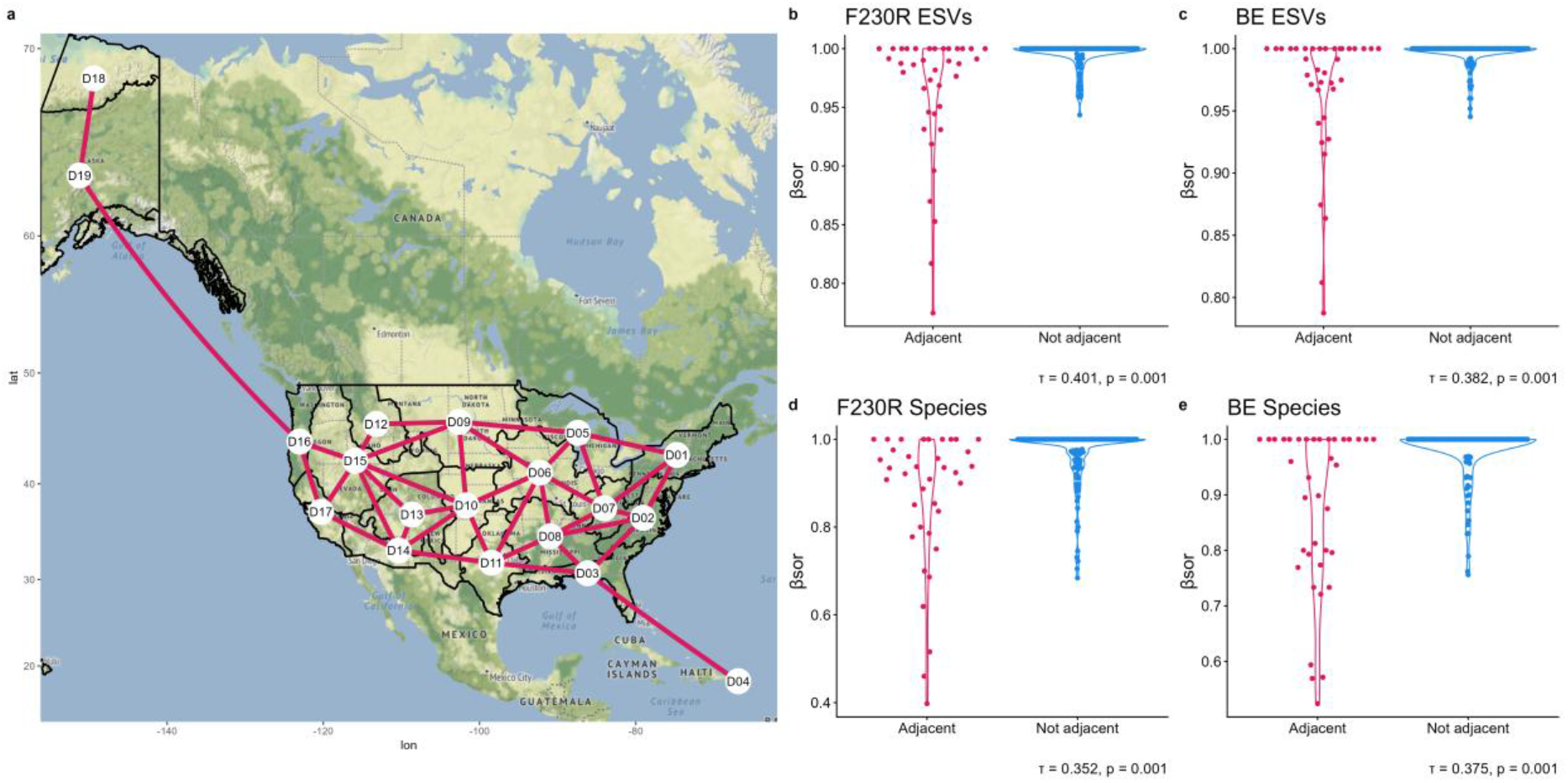
Graph of adjacent ecoregions reveals that β-diversity is spatially autocorrelated for both ESVs and species. The adjacency graph on a map of North America and the ecoregions (a), comparison of β-diversity between adjacent and not adjacent ecoregions for exact sequence variants (ESVs), species, F230R, and BE (b-e). β-diversity was calculated using Sørensen dissimilarity (βsor). The Mantel test was used to assess significance, and correlation was determined using Kendall’s Tau. Kendall’s Tau ranges from -1 to 1, where 1 represent a perfect increasing monotonic relationship and -1 represent a perfect decreasing monotonic relationship. D16 extends into southern Alaska and shares a border with D19, D04 extends into southern Florida and shares a border with D03. Basemap from Stadia Maps https://stadiamaps.com and Open Street Map https://www.openstreetmap.org.

The Sørensen dissimilarity between ecoregions using ESVs and species (where a species is the aggregation of all ESVs within a species bound cluster, SBC, see methods) presence/absence was calculated to represent β-diversity between ecoregions. β-diversity dissimilarity matrices were compared to the adjacency matrix using Kendall rank correlation ^22^. The correlations for both data types and primer sets had p-values as low as possible with the number of permutations performed for the Mantel test (Fig. 1b-e) ^23^. This suggests that ecoregions are spatially autocorrelated, and adjacent ecoregions share more similar biodiversity than ecoregions that are not adjacent.

### B-diversity graphs reveal ecoregion relationships

The correlation strength of our results show that adjacency and β-diversity between ecoregions do not have a perfect agreement. To assess how ecoregions are connected by β-diversity, graphs were created from the dissimilarity matrices. For each graph, the percolation threshold was determined to filter out edges from the graph ^24–26^. Edges with values above the percolation threshold were removed (Fig. 2).

**Fig. 2.**
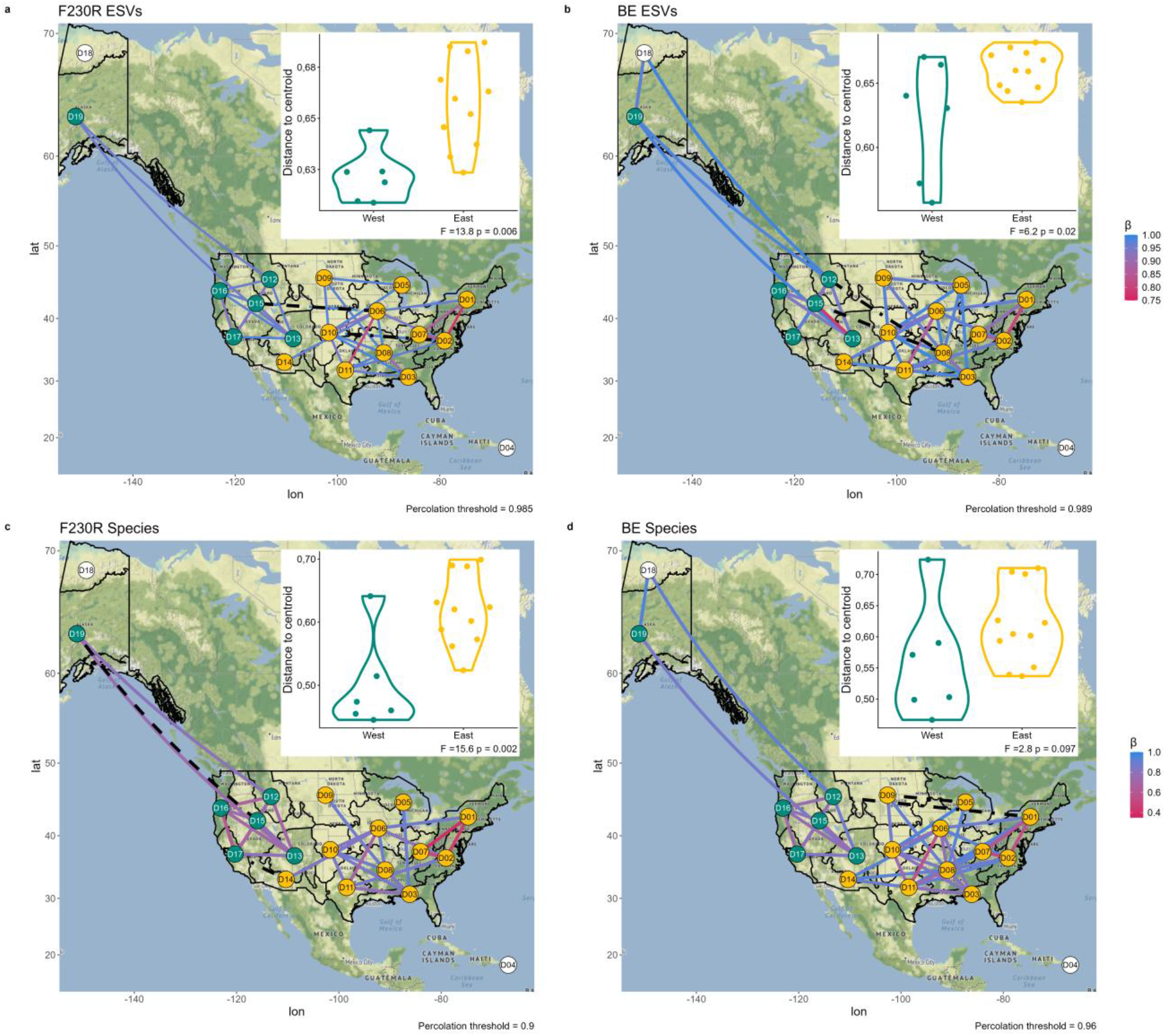
β-diversity graphs separate ecoregions into two groups located in western and eastern United States. Graphs of β-diversity were created for exact sequence variants (ESVs) and species from F230R (a, b) and ESVs and species from BE (c, d) using Sørensen dissimilarity matrices, edges that had values above the percolation threshold were removed from the graph. Inset plots show a comparison of β-diversity dispersion between East and West ecoregions with significance being tested using a Permutational Multivariate Analysis of Variance. Black dot dash lines indicate edges with the highest Sørensen dissimilarity and black dash lines indicate edges with the second highest Sørensen dissimilarity. Basemap from Stadia Maps https://stadiamaps.com and Open Street Map https://www.openstreetmap.org.

For all graphs, the D04 ecoregion was disconnected, and D18 was disconnected from the graphs generated from data generated from F230R amplicons. It was observed that the rest of the ecoregions formed into two groups that were weakly connected within the graphs. These groups were named East and West based on their relative geographic position. Edges in the graph that connected East and West groups had either the closest or second closest β-diversity values to their respective percolation threshold, and none of the edges connecting East and West were consistently observed in any of the four graphs. The dispersion of β-diversity was generally higher in East ecoregions when compared to West ecoregions.

The spatial autocorrelation for the East and West groups was also calculated (Table 1). Interestingly, in the West the correlation at the ESV level was much stronger than at the species level, where the correlation was very weak. In the West, only F230R ESVs had significant spatial autocorrelation. The correlation in the East was always significant.

**Table 1.** Spatial autocorrelation of ecoregion adjacency matrix and β-diversity for species and ESVs within East and West ecoregion groups. Significance was tested using the Mantel test.

|  |  | **West** |  | **East** |  |
|---|---|---|---|---|---|
|  |  | **Kendall's τ** | **p-value** | **Kendall's τ** | **p-value** |
| **F230R** | **ESV** | 0.398 | 0.028 | 0.322 | 0.006 |
|  | **Species** | 0.013 | 0.531 | 0.330 | 0.001 |
| **BE** | **ESV** | 0.363 | 0.089 | 0.305 | 0.017 |
|  | **Species** | 0.112 | 0.389 | 0.373 | 0.002 |

### Intraspecific genetic differentiation

Of the 387 species from F230R that were analyzed, 53 were identified in two or more East ecoregions exclusively, 27 were identified in two or more West ecoregions exclusively, and 8 were identified in both East and West ecoregions. Of the 292 species from BE, 41 species were identified in two or more East ecoregions exclusively, 16 were identified in two or more West ecoregions exclusively, and 1 was identified in both East and West ecoregions.

Species were organized into a neighbour-joining tree based on their ESV with the most reads (Fig. 3). Species generally clustered into their assigned orders, however all Plecoptera did not cluster together for F230R and all Ephemoptera did not cluster together for F230R and BE. The intraspecific genetic differentiation, the number of ESVs, the d_min_ (see species bound clusters in methods for more details), and the number of ecoregion was compared between species exclusively in the East and West groups (Fig. S1). To calculate the intraspecific genetic differentiation, the mean pairwise Sørensen dissimilarity between ecoregions of ESVs within species was calculated. The intraspecific genetic differentiation was significantly higher for species in the West ecoregions when compared to those in the East, and the d_min_ of BE species in the West was also significantly higher.

**Fig. 3.**
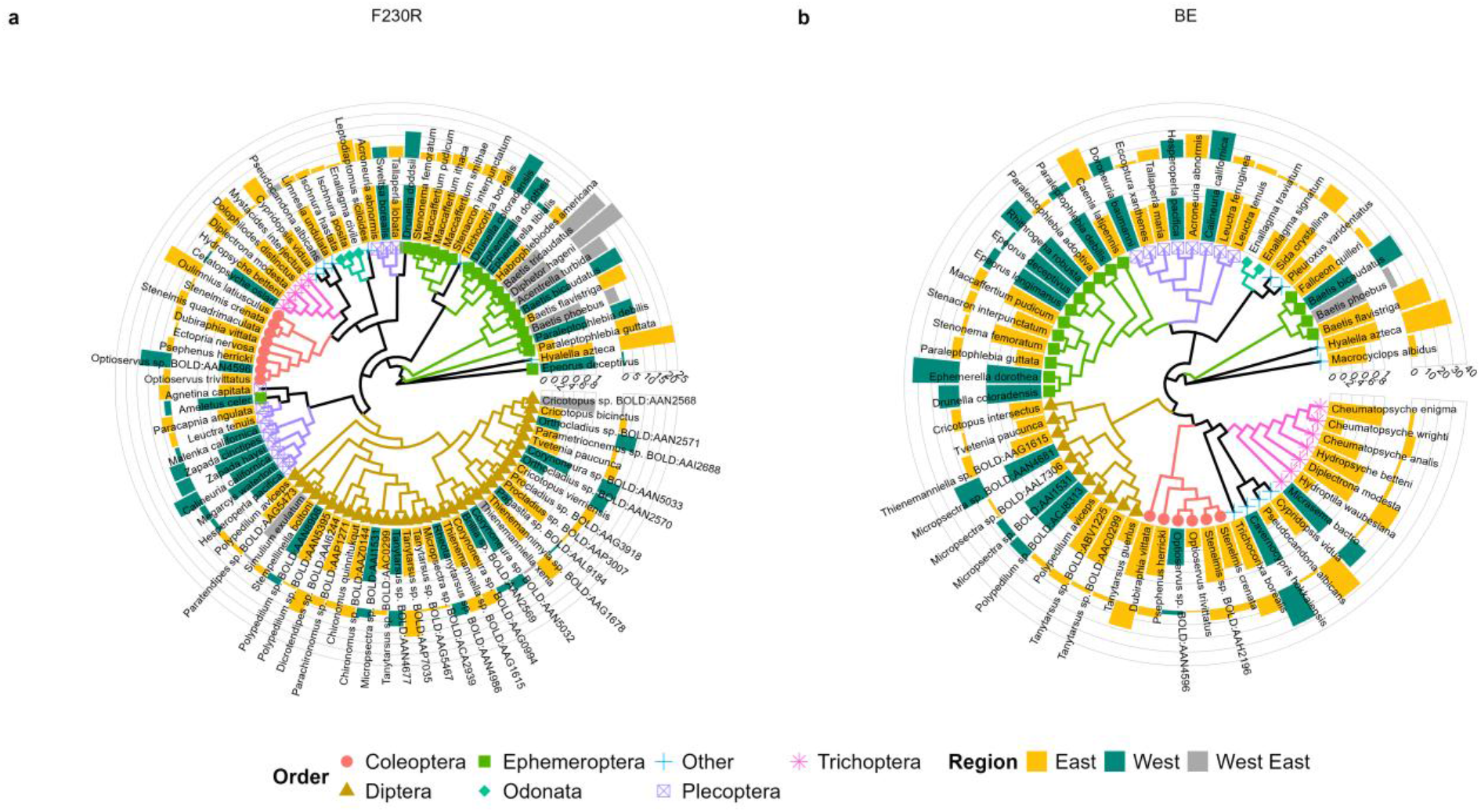
Neighbour-joining trees comparing species in East and West ecoregions. Neighbour-joining trees were created using the ESV sequence with the highest number reads for each species for F230R (a) and BE (b). The inner bar graph is the intraspecific genetic differentiation that was calculated using the mean pairwise Sorensen dissimilarity between ecoregions for each species. The outer bar graph is the d_min_ for each species, where d_min_ is the lowest threshold that each species cluster forms, see species bound clusters in methods for more details. The d_min_ of *Hylalella azteca* was masked by setting it to the next highest d_min_ value, *H. azteca* had a d_min_ of 43 for F230R and a d_min_ of 65 for BE. *H. azteca* has been documented as a species complex ^27^.

## Discussion

In this study we demonstrated that ecoregion structural connectivity and functional connectivity are spatially autocorrelated (Fig. 1). The link between ecoregion adjacency and organism movement can be intuitively understood. For an organism to move from one ecoregion to any other ecoregion, it must always pass through an adjacent ecoregion. Therefore, ecoregions that are adjacent will have more individuals move between them than those that are not adjacent. The agreement between structural and functional connectivity is further exemplified by ecoregions D04 and D18 (for F230R). Structurally, both ecoregions are only adjacent to one ecoregion and functionally, both ecoregions are disconnected from the β-diversity graph above the percolation threshold (Fig. 2). This suggests that these ecoregions are both structurally and functionally isolated.

Adjacency of ecoregions is a simple way of investigating structural connectivity, it only takes into account the spatial arrangement of ecoregions and ignores important features like variations in ecotones and treats the entirety of the ecoregion as homogenous. In the case D04, this ecoregion covers southern Florida as well as the entirety of Puerto Rico, however all sampling sites are on Puerto Rico. This means that all the aquatic arthropods sampled in D04 are separated from all other samples by an ocean. The ocean acts as a barrier to dispersal for our target species, for an organism to cross the ocean, it either needs strong dispersal capabilities or resilience to a marine environment ^28^. If sites were sampled in southern Florida, functional isolation of D04 may be less exaggerated.

Considering where structural and functional connectivity diverge can provide insight into how ecotones may be restricting connectivity. From β-diversity graphs (Fig. 2), there is a clear separation of western and eastern ecoregions. The ecotones where East meets West corresponds roughly with the abrupt and distinct boundary of the Great Plains and the Rocky Mountains ^29^. Mountains are well documented barriers to dispersal that will restrict the movement of organisms and gene flow ^30–33^. It is likely the transition between plains and mountains that is resulting in the increase of β-diversity between these adjacent ecoregions.

Within the eastern ecoregions, there is greater β-diversity dispersion than in the western ecoregions (Fig. 2). While this could indicate there is greater variation in organism composition in the East, it is likely a reflection of its larger size ^34^. Conversely, intraspecific genetic differentiation in the West ecoregions was significantly higher than in the East ecoregions (Fig. 3 and S1). This suggests that the gene flow for species in the western United States is more restricted than in the eastern United States. The restriction of gene flow in the western ecoregions could corresponds with the Rocky Mountains acting as sky islands. Studies of butterflies in the Rocky Mountains by DeChain and Martin suggested that populations underwent periods of expansion and contraction ^35–37^. During times of glaciation, cooler climates would connect areas of higher elevation allowing for expansion, while during interglaciation, warmer climates would isolate populations on mountains. We are currently in a period of warming climates which is likely isolating montane populations and restricting gene flow. This pattern of expansion and contraction could also explain why ESVs have higher spatial autocorrelation than species in the western ecoregions (Table 1). Some ESVs, which represent a more recent timescale, could be undergoing expansion during the warming climate, while species, representing a longer timescale, are the reflection of previous climatic changes that resulted in isolation.

Isolation and connectivity are key opposing factors in species persistence. If two populations become isolated from each other, gene flow between these populations will cease. If both populations are of a sustainable size, changes in genetic structure can occur through adaptations and genetic drift, eventually resulting in speciation. This is an ideal outcome of isolation from a conservation perspective ^38^. However, if an isolated population’s size is (or becomes) too small, genetic drift can lead to a loss of genetic diversity weakening the populations ability to adapt and inbreeding depression can lead to the accumulation of deleterious mutations ^39,40^. These factors will reduce the population’s fitness and can push it to extinction ^41^.

An important consideration from our study is the role of anthropogenic stressors in influencing the dynamics of change in populations with different degrees of connectivity. Connectivity measurements have been incorporated in policies and plans for conservation of nature ^42^. Fine scale gene flow analysis among populations of multiple species provide a unique biological data layer to map functional connectivity at large-scale. Our observation that species have higher intraspecific genetic differentiation in the West suggests that these species are more isolated, and therefore could have slower recovery from anthropogenic stressors through connected populations. Incorporating a metabarcoding-based analysis of genetic diversity can significantly aid our efforts to mitigate the impacts of climate change and biodiversity loss, by providing insights linked to genetic structure of many species in an ecosystem, leading to better environmental and conservation decision-making.

However, the changes that DeChaine and Martin observed in butterfly species are estimated to have occurred between 40 and 200 thousand years ago ^37^. While more recent than the formation of the Rocky Mountains (millions of years), this timescale is much older than the anthropogenic stressors we are concerned about today. Further incorporating time into the analysis of DNA metabarcoding will be essential to understand how biodiversity is dynamically changing under these pressures ^17^. Publicly accessible data from long term projects like NEON that are continuing to generate new data over multiple years will be essential to combat biodiversity loss.

## Conclusion

We analyzed the connectivity of DNA metabarcoding data from benthic arthropods collected across the United States. We found the adjacency of geo-clustered ecoregions (structure) was correlated with the β-diversity (function) of ESVs and species. However, adjacent ecoregions along the border of the Rocky Mountains had high β-diversity. This allowed ecoregions to be separated into East and West groups. Species in the western group had higher intraspecific genetic diversity, likely corresponding to the heterogeneity in the Rocky Mountains. This could indicate that these species are more isolated and therefore more at risk of anthropogenic stressors due to a lack of connectivity. DNA metabarcoding has multiple layers of information from many taxa that can be used to better understand connectivity and improve conservation planning.

## Methods

### Data set

Macroinvertebrate metabarcoding data was download from the National Ecological Observatory Network on September 29^th^ 2025 ^43^. Raw sequence files were downloaded using wget and urls in rawDataFilePath. The data set covered 19 ecoregions, 34 sites, and 470 samples that were collected from 2018 to 2022. Not every site was sampled every year, however every site was sampled in at least two years. Sites included three different systems: lakes, rivers, and streams. The sampling protocol is available from NEON in AOS PROTOCOL AND PROCEDURE: INV – AQUATIC MACROINVERTEBRATE SAMPLING. The sample processing protocol is available from NEON in Laragen Invertebrate Metabarcode SOP: NEON Invertebrate Metabarcoding. During sample processing two amplicons were targeted during PCR using the F230R primers and the BE primers ^44–46^. An Illumina Miseq with a v2 500-cycle kit (cat# MS-1022003) was used to sequence amplicons.

### DNA metabarcoding processing

Raw sequence files were processed using the same steps as Riley *et al*. ^33^. Briefly, the MetaWorks 1.13.0 pipeline was run using the default ESV and COI settings ^47^. Within the pipleline, SEQPREP v1.3.2 merged raw paired-end reads ^48^, CUTADAPT v4.1 trimmed primers ^49^, and VSEARCH v2.21.1 dereplicated, denoised (*α* = 2.0, minimum reads = 8), and removed putative chimeric sequences ^50^. ESVs were assigned to taxonomy using the using the Ribosomal Database Project (RDP) classifier for Eukaryote COI mtDNA sequences (RDP COI v5.1.0) ^51,52^. Only ESVs that were assigned to arthropods were processed further. To remove putative pseudogenes, the profile hidden Markov model (HMM) of ESVs translated using ORFinder ^53^ were compared to a profile HMM of BOLD arthropod COI barcode sequences ^54^ using HMMER v3.3.2 ^55^. ESVs with sequence bit scores below the 25^th^ percentile were identified as outliers and removed. After running MetaWorks, data was processed using Python 3.9.2 ^56^. An additional multiple sequence alignment filtering step was performed using MAFFT to ensure all sequences had the same length while removing sequences that contain putative insertions/deletions ^57^. Finally, ESVs with fewer than 100 reads were filtered out as an additional quality control step.

### Species bound clusters (SBCs)

ESVs were clustered into species bound clusters (SBCs), which are intended to represent species ^33^. They are defined by the taxonomic assignment of ESVs and sequence similarity. ESVs with a species level bootstrap value >= 0.8 from the RDP classifier were assigned to a species. Using SWARM 3.1.3, ESVs are clustered from d = 0 (clustering threshold, where d is the number of nucleotide differences between sequences) until the maximum d value where all ESVs belong to the same cluster ^58^. In SWARM the –n option is used, this option stops clusters from being broken based on ESV abundance, which can result in the clusters losing their nested structure. Within the full set of nested clusters, the smallest clusters that include all ESVs assigned to one species and no ESVs assigned to a different species are identified, these clusters are SBCs. The boundary of a cluster is defined by taxonomic assignment. The d_min_ value represents the lowest threshold that the cluster is formed, the d_max_ value represents the highest threshold that the cluster exists.

### Analysis

R 4.2.1 was used to execute all remaining analyses ^59^. Two additional filtering steps were performed to avoid cross contamination of samples. First, ESVs with fewer than 5 reads were filtered out samples to avoid tag-switching between samples. Second, every site included in this data set was sampled at least twice between 2018 and 2022. In this study we focused on studying patterns in space and not time. Therefore, time could be used as a filter where ESVs and species that were only detected in an ecoregion in one year were removed. This was done at the risk of removing rare and vagrant ESVs and species, since it is unlikely the same ESVs/species would cross contaminate samples in the same ecoregions over multiple years.

All samples (replicates, sites, years) were pooled within each ecoregion and reads counts were converted into presence/absence data. This data is unlikely to represent sampling effort in each ecoregion but should be considered as a list of everything that has ever been observed in each ecoregion. B-diversity between ecoregions was calculated using beta.pair.abund with output beta.bray (equivalent to beta.pair function and beta.sor output for presence/absence data) in the R package betapart ^60^. The mantel function from the vegan R package with method=“kendall” was used to calculate the correlation and significance between β-diversity matrices and the adjacency matrix ^61^. The perc.thr function from the sidier R package was used to calculate the percolation threshold, percolation thresholds were tested increasing by 0.001 over a range of 0 to 1 ^25^. Over this range of tested values, the percolation threshold is determined as the point where the graph separates into its dominant clusters. The betadisper function from the vegan R package was used to calculate dispersion and dispersion between groups was compared using the adonis2 function from the vegan R package ^61^. To create neighbour-joining trees, the dist.alignment function the seqinr R package was used to create distance matrix and the bionj function from the ape R package was used to turn this distance matrix into a tree ^62,63^. Finally, the wilicox.test function from the stats R package was used to compare the mean pairwise Sørensen dissimilarity, the number of ESVs, the d_min_, and the number of ecoregion between groups. Statistical test were considered significant if p-values were less than 0.05.

The following R packages were also used for data manipulation: dplyr, reshape2, and stringr ^64–66^. The following R packages were also used in figure generation: cowplot, igraph, ggbeeswarm, ggmap, ggnetwork, ggnew scale, ggplot2, ggpubr, ggtree, ggtreeExtra, rgdal, and sf^67–83^.

## Data availability

Raw sequence files are available from the National Ecological Observatory Network under the product ID DP1.20126.001 ^43^.

## Code availability

Processed data and code for analysis and figure generation is available at https://github.com/anriley/Connectivity-and-genetic-diversity.

## Author contributions

Conceptualization: ACR, MH

Methodology: ACR, MH

Investigation: ACR

Visualization: ACR, MH

Funding acquisition: MH

Writing – original draft: ACR, MH

Writing – review & editing: ACR, MH

## Supplementary Materials

Figs. S1

