## Supplemental Figure S1 for "Continental scale genetic diversity supports geo-clustered ecoregions connectivity"

A. C. Riley, M. Hajibabaei

**The file includes:**

Figs. S1

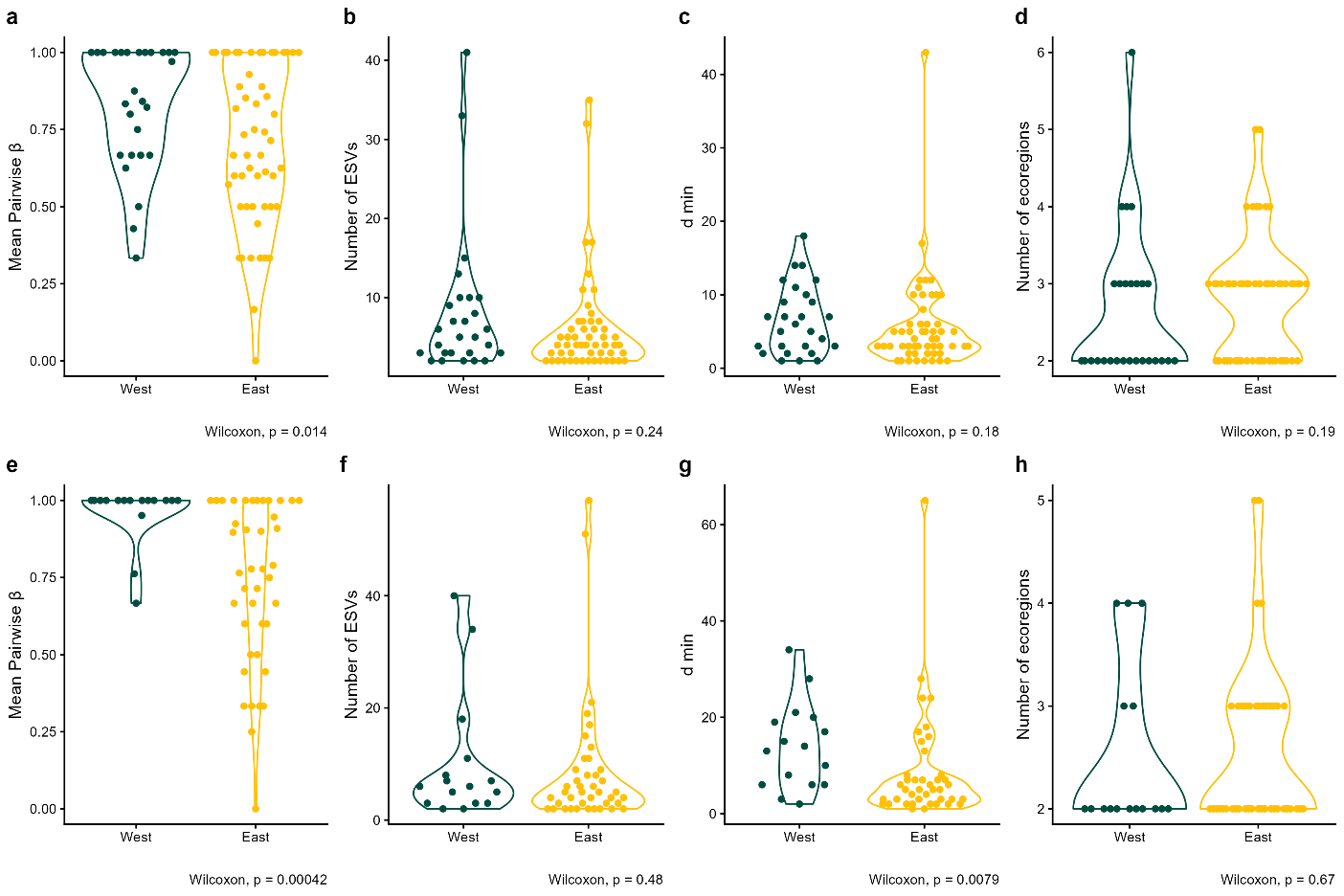

Fig. S1.

Comparison of species traits in East and West ecoregions. Comparisons were made for F230R (a-d) and BE (e-h). Intraspecific genetic differentiation (a, e), number of exact sequence variants (ESVs) (b, f), d_min_ (c, g), and number of ecoregions (d, h) for each species was compared.
